# MicroRNAs from the parasitic plant *Cuscuta campestris* target host messenger RNAs

**DOI:** 10.1101/180497

**Authors:** Saima Shahid, Gunjune Kim, Nathan R. Johnson, Eric Wafula, Feng Wang, Ceyda Coruh, Vivian Bernal-Galeano, Tamia Phifer, Claude W. dePamphilis, James H. Westwood, Michael J. Axtell

## Abstract

Dodders (*Cuscuta* spp.) are obligate parasitic plants that obtain water and nutrients from the stems of host plants via specialized feeding structures called haustoria. Dodder haustoria facilitate bi-directional movement of viruses, proteins, and mRNAs between host and parasite^1^, but the functional effects of these movements are not clear. Here we show that *C. campestris* haustoria accumulate high levels of many novel microRNAs (miRNAs) while parasitizing *Arabidopsis thaliana* hosts. Many of these miRNAs are 22 nts long, a usually rare size of plant miRNA associated with amplification of target silencing through secondary small interfering RNA (siRNA) production^2^. Several *A. thaliana* mRNAs are targeted by *C. campestris* 22 nt miRNAs during parasitism, resulting in mRNA cleavage, secondary siRNA production, and decreased mRNA accumulation levels. Hosts with mutations in two of the targets supported significantly higher growth of *C. campestris.* Homologs of target mRNAs from diverse plants also have predicted target sites to induced *C. campestris* miRNAs, and the same miRNAs are expressed and active against host targets when *C. campestris* parasitizes a different host, *Nicotiana benthamiana*. These data show that *C. campestris* miRNAs act as *trans*-species regulators of host gene expression, and suggest that they may act as virulence factors during parasitism.

---

Host-induced gene silencing (HIGS) involves plant transgenes that produce siRNAs which can silence targeted pathogen/parasite mRNAs in *trans*^3,4^. Plant-based HIGS is effective against fungi^5^, nematodes^6^, insects^7^, and the parasitic plant *Cuscuta pentagona*^8^. The apparent ease of engineering HIGS suggests that plants might exchange naturally occurring small RNAs during pathogen/parasite interactions. Consistent with this hypothesis, small RNAs from the plant pathogenic fungus *Botrytis cinerea* target host mRNAs during infection^9^, and HIGS targeting *B. cinerea Dicer-Like* mRNAs reduces pathogen virulence^10^. Conversely, the conserved miRNAs miR159 and miR166 can be exported from cotton into the fungal pathogen *Verticillium dahliae* where they target fungal mRNAs encoding virulence factors^11^. However, to date, no examples of naturally occurring *trans*-species miRNAs have been described for plant-to-plant interactions.

*Cuscuta* haustoria facilitate bi-directional movement of viruses, proteins, and mRNAs^1^, but the functional effects of these movements are unclear. *Cuscuta* is susceptible to HIGS, so we hypothesized that naturally occurring small RNAs might be exchanged across the *C. campestris* haustorium and affect gene expression in the recipient species. To test this hypothesis, we profiled small RNA expression from *C. campestris* grown on *A. thaliana* hosts using high-throughput small RNA sequencing (sRNA-seq). Two biological replicates each from three tissues were analyzed: Parasite stem (PS), comprising a section of *C. campestris* stem above the site of haustorium formation; Interface (I), comprising *C. campestris* stem with haustoria with associated *A. thaliana* stem tissue; and Host stem (HS), comprising sections of *A. thaliana* stems above the interface region, as previously described^12^. Small RNA-producing loci from both organisms were identified, classified, and subject to differential expression analyses (Supplementary Data 1).

As expected due to dilution of parasite RNA with host RNA, *C. campestris* small RNA loci were generally down-regulated in I relative to PS (Figure 1A). However, 76 *C. campestris* small RNA loci were significantly (FDR <= 0.05) higher in I relative to PS. 43 of these (57%) were *MIRNA* loci as determined by canonical accumulation of a discrete miRNA/miRNA* pair from predicted stem-loop precursors (Figure 1B, Supplementary Data 2-4). RNA blots confirmed I-specific expression (Figure 1C). One of the 43 *MIRNA*s is a member of the conserved *MIR164* family; the other 42 up-regulated *MIRNA*s have no obvious sequence similarity to previously annotated *MIRNA* loci, and none of their mature miRNAs or miRNA*s were perfectly alignable to the *A. thaliana* genome (Supplementary Data 5). Several of the key *MIRNA* loci were detected by PCR of *C. campestris* genomic DNA prepared from four-day old seedlings that had never interacted with a host plant (Extended Data Figure 1). The majority of the induced *C. campestris MIRNA* loci (26/43) produced a 22 nt mature miRNA. 22 nt plant miRNAs are usually less frequent than 21nt miRNAs, and they are strongly associated with secondary siRNA accumulation from their targets^13,14^. Secondary siRNAs are thought to amplify the strength of miRNA-directed gene silencing^2^.

**Figure 1.**
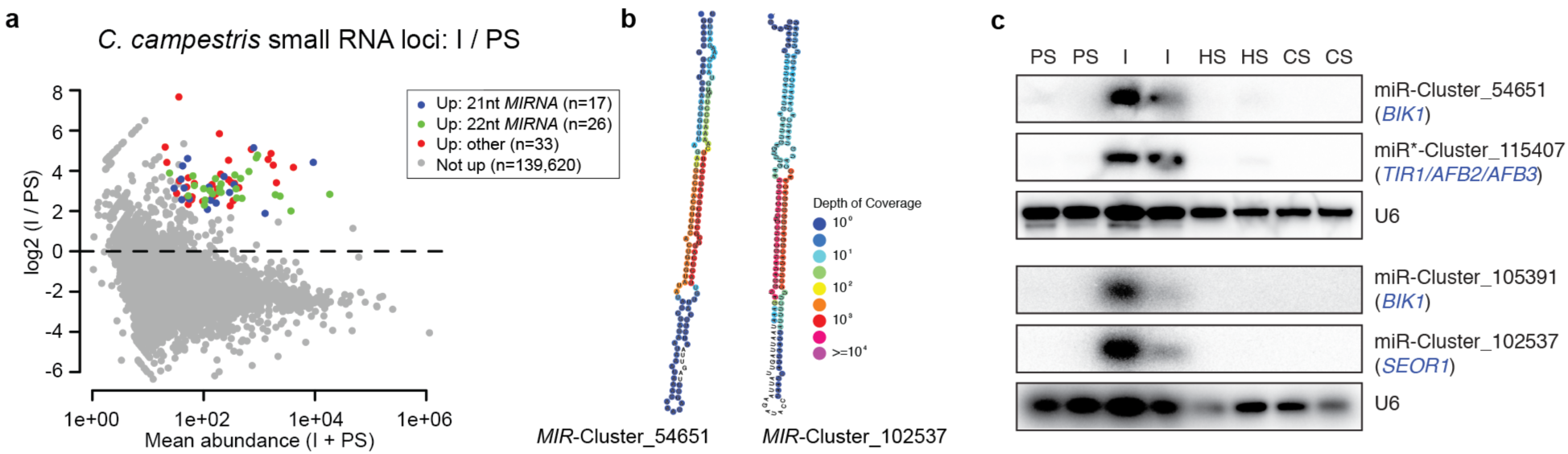
*C. campestris* miRNAs induced at the haustorial interface. **a)** MA plot of *C. campestris* small RNA loci comparing interface (I) to parasite stem (PS) samples. Significantly up-regulated loci (alternative hypothesis: true difference > 2-fold, FDR <= 0.05) are highlighted. **b)** Predicted secondary structures of example *MIRNA* hairpin precursors of *C. campestris* 22 nt miRNAs. Color-coding shows depth of small RNA-seq coverage at each nucleotide. **c)** RNA blots of 22 nt I-induced miRNAs. PS: parasite stem, I: interface, HS: host stem, CS: control stem.

We hypothesized that the induced 22 nt miRNAs would cause secondary siRNA formation from targeted host mRNAs. Therefore we searched for *A. thaliana* mRNAs that had one or more plausible miRNA complementary sites and accumulation of secondary siRNAs specifically in the I small RNA-seq samples. Six *A. thaliana* mRNAs that met both criteria were found: *TIR1*, *AFB2*, and *AFB3*, which encode related and partially redundant auxin receptors^15^, *BIK1*, which encodes a plasma membrane-localized kinase required for both pathogen-induced and developmental signaling^16,17^, *SEOR1*, which encodes a major phloem protein that accumulates in filamentous networks in sieve tube elements and reduces photosynthate loss from the phloem upon injury^18,19^, and *SCZ/HSFB4*, which encodes a predicted transcriptional repressor that is required for the formation of ground tissue stem cells in roots^20^-^22^. The induced siRNAs from these mRNAs resembled other examples of secondary siRNAs in their size distributions, double-stranded accumulation, and phasing (Figure 2A-B; Extended Data Figure 2). *TIR1*, *AFB2*, and *AFB3*, are also known to be targeted by the 22 nt miR393 and to produce secondary siRNAs downstream of the miR393 complementary site^23^. In parasitized stems the location and phase register of the *TIR1*, *AFB2*, and *AFB3* secondary siRNAs shift upstream, proximal to the complementary sites to the *C. campestris* miRNAs (Extended Data Figure 2), implying that the *C. campestris* miRNAs, not miR393, are triggering the I-specific secondary siRNAs. The predominant 21 nt phase register at several loci was shifted +1 to +2 relative to the predictions. This is consistent with the 'phase drift' seen at other phased siRNA loci^24,25^ and likely due to the presence of low levels of 22nt siRNAs, causing the register to be shifted forward. Analysis of uncapped mRNA fragments using 5'-RNA ligase-mediated rapid amplification of cDNA ends found strong evidence for miRNA-directed cleavage at all of the complementary sites to *C. campestris* miRNAs, specifically from interface samples but not from control stem samples (Figure 2; Extended Data Figure 2). We did not find any induced miRNAs or siRNAs from the *A. thaliana* host capable of targeting these six mRNAs. We also did not find any endogenous *C. campestris* secondary siRNA loci corresponding to any of the induced miRNAs. Some *C. campestris* orthologs of *TIR/AFB, SCZ/HSFB4*, and *BIK1* had possible, but very poorly complementary, miRNA target sites (Extended Data Figure 3). These observations suggest that the induced *C. campestris* miRNAs have evolved to avoid targeting ‘self’ transcripts. We conclude that 22 nt miRNAs from *C. campestris* act in a *trans*-species manner to target *A. thaliana* mRNAs.

**Figure 2.**
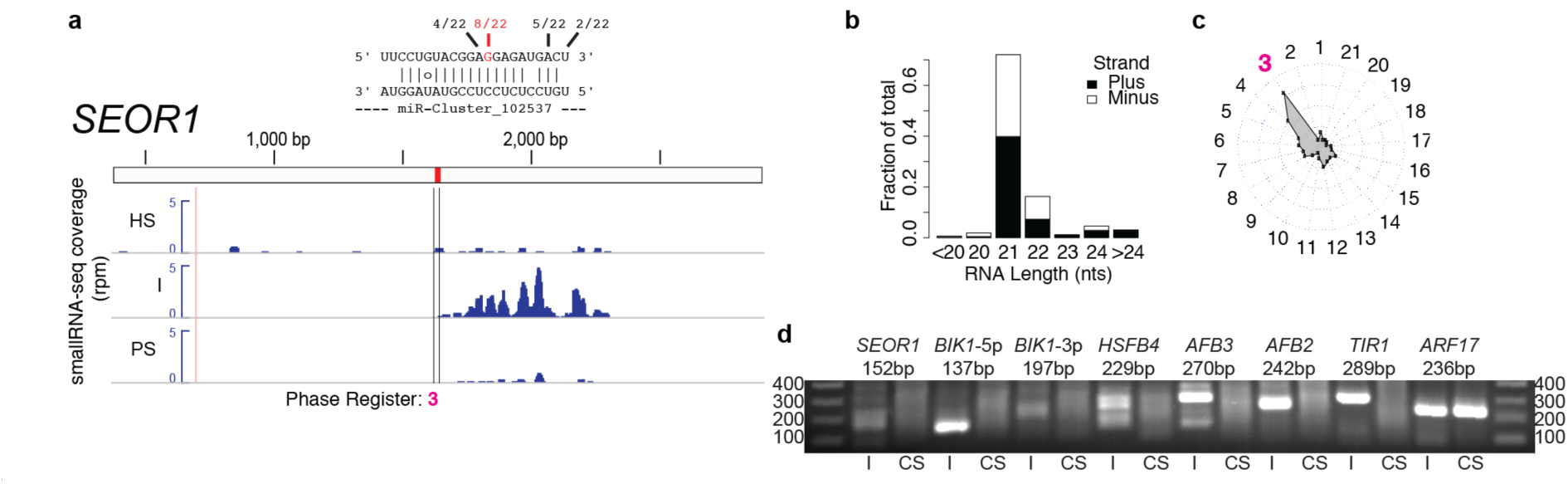
*C. campestris* miRNAs cause slicing and phased siRNA production from host mRNAs. **a)** Small RNA-seq coverage (reads per million) for the *A. thaliana SEOR1* transcript for host stem (HS), interface (I), and parasite stem (PS) samples. miRNA complementary site with 5'-RLM-RACE data and expected 21nt phasing register is shown. **b)** Length and polarity distribution of *SEOR1*-mapped siRNAs from the I samples. **c)**. Radar chart showing fraction of I-derived siRNAs in each of the 21 possible phasing registers. **d)** 5'-RLM-RACE products from nested amplifications for the indicated cDNAs. I: interface, CS: control stem. *ARF17* is a positive control.

**Figure 3.**
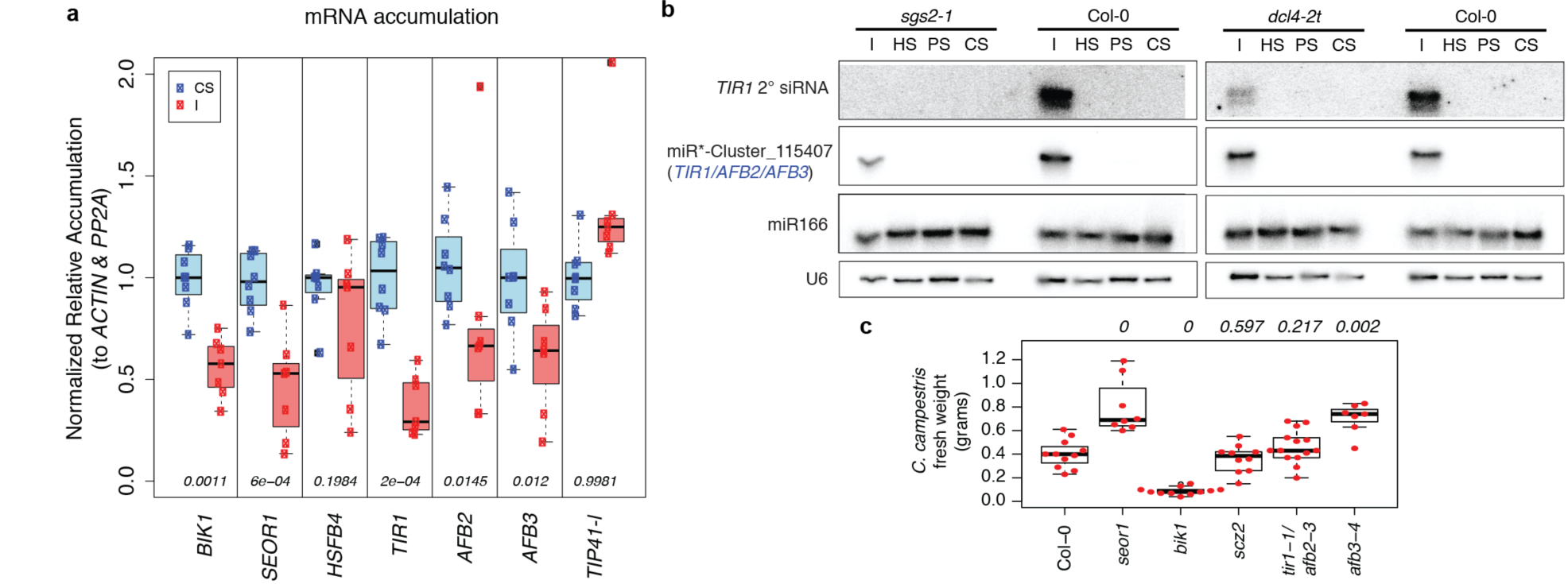
Effects of *C. campestris* miRNAs and their targets. **a)** *A. thaliana* mRNA accumulation levels in I (interface) vs. CS (control stems) during *C. campestris* parasitism, assessed by qRT-PCR. Data from 7 or 8 biological replicates are plotted (dots), and boxplots indicate the median (horizontal lines), 1st-3rd quartile range (boxes), and up to 1.5 x the interquartile range (whiskers). Numbers indicate p-values comparing CS and I samples (Wilcoxon rank-sum test, unpaired, one-tailed). *TIP41-l*: control mRNA. **b)** RNA blots from *C. campestris* infestations of the indicated *A. thaliana* genotypes. I: interface, HS: host stem, PS: parasite stem, CS: control stem. **c)** Accumulation of *C. campestris* biomass on *A. thaliana* hosts of the indicated genotypes 18 days post-attachment. P-values (Wilcoxon rank-sum tests, unpaired, two-tailed) from comparison of mutant to wild-type (Col-0) are shown. Boxplot conventions as in panel a. n=11, 8, 11, 10, 14, and 7 for Col-0, *seor1*, *bik1, scz2, tir1-1/afb2-3*, and *afb3*, respectively.

Accumulation of five of the six secondary siRNA-producing targets was significantly reduced in stems parasitized by *C. campestris* compared to un-parasitized stems (Figure 3A), consistent with miRNA-mediated repression. The true magnitude of repression for these targets could be even greater, since many miRNAs also direct translational repression. Accumulation of many known *A. thaliana* secondary siRNAs is often partially dependent on the endonuclease Dicer-Like 4 (DCL4) and wholly dependent on RNA-Dependent RNA polymerase 6 (RDR6/SGS2/SDE1)^2^. Accumulation of an abundant secondary siRNA from *TIR1* was eliminated entirely in the *sgs2-1* mutant, and reduced in the *dcl4-2t* mutant (Figure 3B). Thus host *DCL4* and *RDR6/SGS2/SDE1* are required for secondary siRNA production. This implies that the *C. campestris* derived miRNAs are active inside of host cells and hijack the host’s own silencing machinery to produce secondary siRNAs.

In repeated trials with varying methodologies we did not observe consistent significant differences in parasite fresh weight using *dcl4-2t* and *sgs2-1* mutants as hosts (Extended Data Figure 4). Thus, loss of induced secondary siRNAs is not sufficient to detectably affect parasite biomass accumulation in this assay. Similarly, there were no significant differences in parasite fresh weights when *scz2* or *tir1-1/afb2-3* plants were used as hosts (Figure 3C). Significantly less *C. campestris* biomass was observed using *bik1* mutants as hosts. However, interpretation of this result is complicated by the weak, frequently lodging stems of the *bik1* mutant^16^. Significantly more *C. campestris* biomass was observed when grown on *seor1* or *afb3-4* mutant hosts (Figure 3C). Therefore, both *SEOR1* and *AFB3* function to restrict *C. campestris* growth on *A. thaliana*. This observation is consistent with the hypothesis that both *SEOR1* and *AFB2* are *trans*-species miRNA targets of biological relevance.

**Figure 4.**
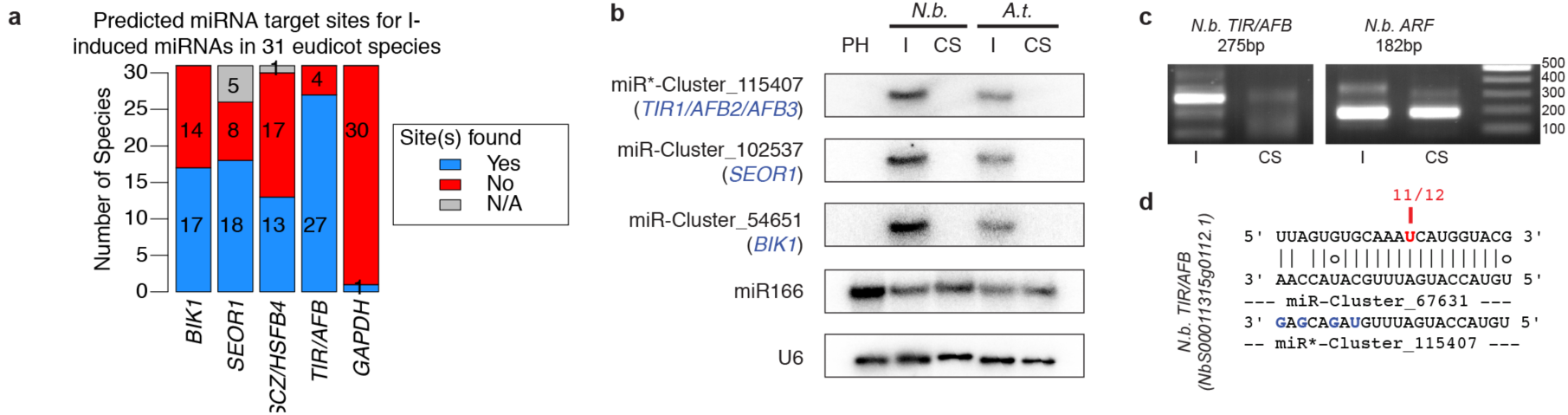
Conservation of host mRNA targeting by *C. campestris*. **a)** Predicted targets of the induced *C. campestris* miRNAs/miRNA*s among indicated orthologs. **b)** RNA blots from interface (I) and control stem (CS) samples of *C. campestris* infested *N. benthamiana* (*N.b.*) and *A. thaliana* (*A.t.*) hosts, as well as *C. campestris* pre-haustoria (PH; Extended Data Figure 8). **c)** 5'-RLM-RACE products for the indicated *N. benthamiana* cDNAs. I: interface, CS: control stem. *N.b. ARF* is a positive control. Image cropped to remove irrelevant lanes; uncropped image: Extended Data Figure 7. **d)** Complementary site and 5'-RLM-RACE results from a *N. benthamiana TIR/AFB* ortholog.

*C. campestris* has a broad host range among eudicots^26^. Therefore, we searched for miRNA complementary sites for the interface-induced *C. campestris* miRNAs in eudicot orthologs of the targeted *A. thaliana* mRNAs. Probable orthologs of *BIK1, SEOR1, TIR/AFB*, and *SCZ/HSFB4* were predicted targets of interface-induced miRNAs in many eudicot species, while only one species had predicted targets for the negative control orthologs of *GAPDH* (Figure 4A, Extended Data Table 1). We conclude that the induced *C. campestris* miRNAs collectively would be able to target *TIR/AFB, SEOR1, SCZ/HSFB4*, and *BIK1* orthologs in many eudicot species.

We performed additional small RNA-seq from *C. campestris* on *A. thaliana* hosts, and from *C. campestris* on *Nicotiana benthamiana* hosts. Both sets of experiments were designed identically to the original small RNA-seq study (two biological replicates each of HS, I, and PS samples). The I-induced set of *C. campestris MIRNA* loci was highly reproducible across both of the *A. thaliana* experiments as well as the *N. benthamiana* experiment (Extended Data Figure 5). Induction of several *C. campestris* miRNAs during *N. benthamiana* parasitism was confirmed by RNA blots (Figure 4B). Several *N. benthamiana* mRNAs had both plausible target sites for *C. campestris* miRNAs and accumulation of phased, secondary siRNAs in the I samples, including orthologs of *TIR/AFB* and *BIK1* (Extended Data Figure 6). Analysis of uncapped RNA ends provided strong evidence for miRNA-directed cleavage of one of the *N. benthamiana TIR1/AFB* orthologs (Figures 4C-D; Extended Data Figure 7). This directly demonstrates that the same *C. campestris* miRNAs target orthologous host mRNAs in multiple species. None of the interface-induced miRNAs we tested were detectable from *C. campestris* pre-haustoria sampled from seedling tips that had coiled around dead bamboo stakes instead of a live host (Figure 4B; Extended Data Figure 8). This suggests that expression of these miRNAs requires prior contact with a living host.

These data demonstrate that *C. campestris* induces a large number of miRNAs at the haustorium, and that some of them target host mRNAs and reduce their accumulation. Many of the induced miRNAs are 22 nts long, and associated with secondary siRNA production from their host targets using the host’s own secondary siRNA machinery. Several of the targets are linked to plant pathogenesis: Manipulation of *TIR1/AFB2/AFB3* accumulation levels affects bacterial pathogenesis and defense signaling^27^, and BIK1 is a central regulator of pathogen-induced signaling^28^. Perhaps the most intriguing target is *SEOR1*, which encodes a very abundant protein present in large agglomerations in phloem sieve-tube elements^18^. *seor1* mutants have an increased loss of sugars from detached leaves^19^, and our data show that *seor1* mutants also support increased *C. campestris* biomass accumulation. A key function of the haustorium is to take nutrients from the host phloem; targeting *SEOR1* could be a strategy to increase sugar uptake from host phloem. Overall, these data suggest that *C. campestris trans*-species miRNAs might function as virulence factors to remodel host gene expression to its advantage during parasitism.

**Methods** [separate online only document].

## Acknowledgements

We thank the Penn State & Huck Institutes Genomics Core Facility for small RNA-seq services. We thank Hervé Vaucheret, Michael Knoblauch, Gabriele Monshausen, Tesfaye Mengiste, and Renze Heidstra for gifts of *A. thaliana* mutant seed. We thank Beth Johnson for advice on growing conditions for *C. campestris*. Purchase of the Illumina HiSeq2500 used for small RNA-seq was funded by a major research instrumentation award from the US National Science Foundation [1229046 to MJA and CWD]. This research was supported in part by awards from the US National Science Foundation [1238057 to JHW and CWD; 1339207 to MJA] and the National Institute of Food and Agriculture [135997 to JHW].

## Author Contributions

SS and MJA performed most bioinformatics analysis. SS, MJA, and NRJ prepared figures and tables. GK and JHW cultivated and harvested plant specimens used for initial small RNA-seq experiments. NRJ, SS, TP, and MJA cultivated and harvested plant specimens for other experiments. EW, GK, CWD, and JHW performed genome and transcriptome sequencing and preliminary assemblies. FW, SS, and NRJ performed RNA blots. SS and MJA performed 5'-RLM-RACE and qRT-PCR. CC and TP constructed small RNA-seq libraries. NRJ and VB-G performed growth assays. MJA and JHW conceived of the project. MJA wrote and revised the manuscript with input from all other authors.

## Author Information

The authors declare no competing financial interests

## Extended Data

**Extended Data Figure 1.**
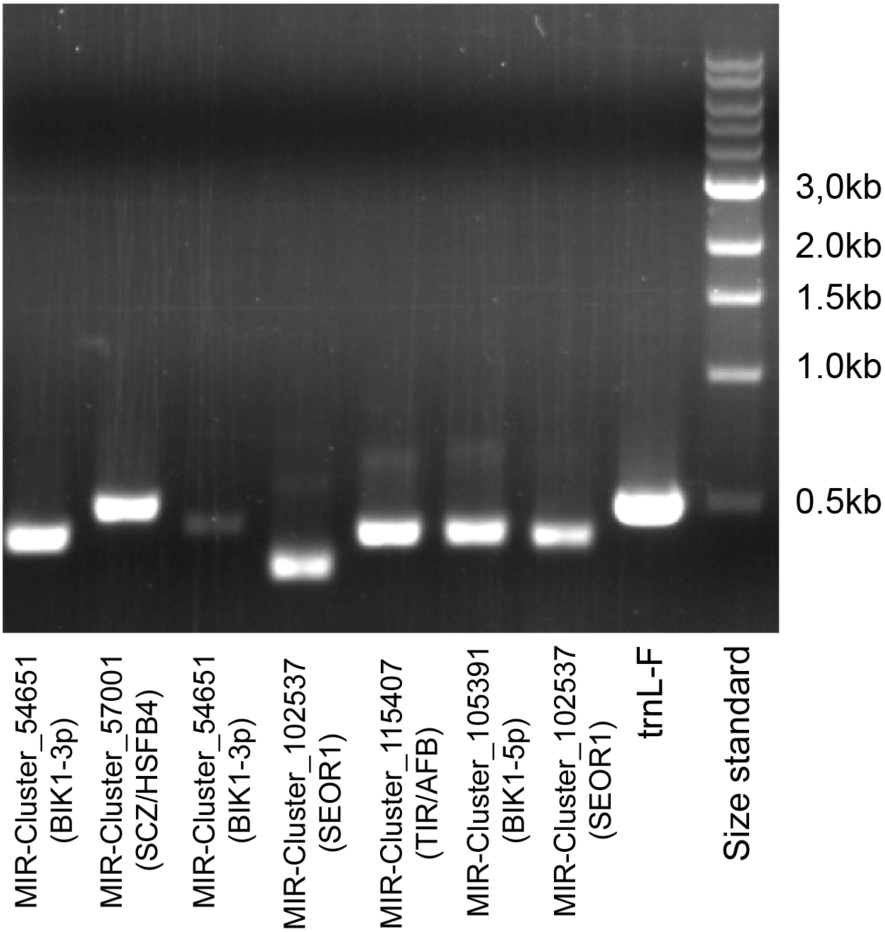
PCR of *C. campestris MIRNA* loci. The template for PCR was genomic DNA isolated from *C. campestris* seedlings four days after germination; the seedlings had never attached to nor been near a host plant, ruling out host DNA contamination. trnL-F: Positive control plastid locus.

**Extended Data Figure 2.**
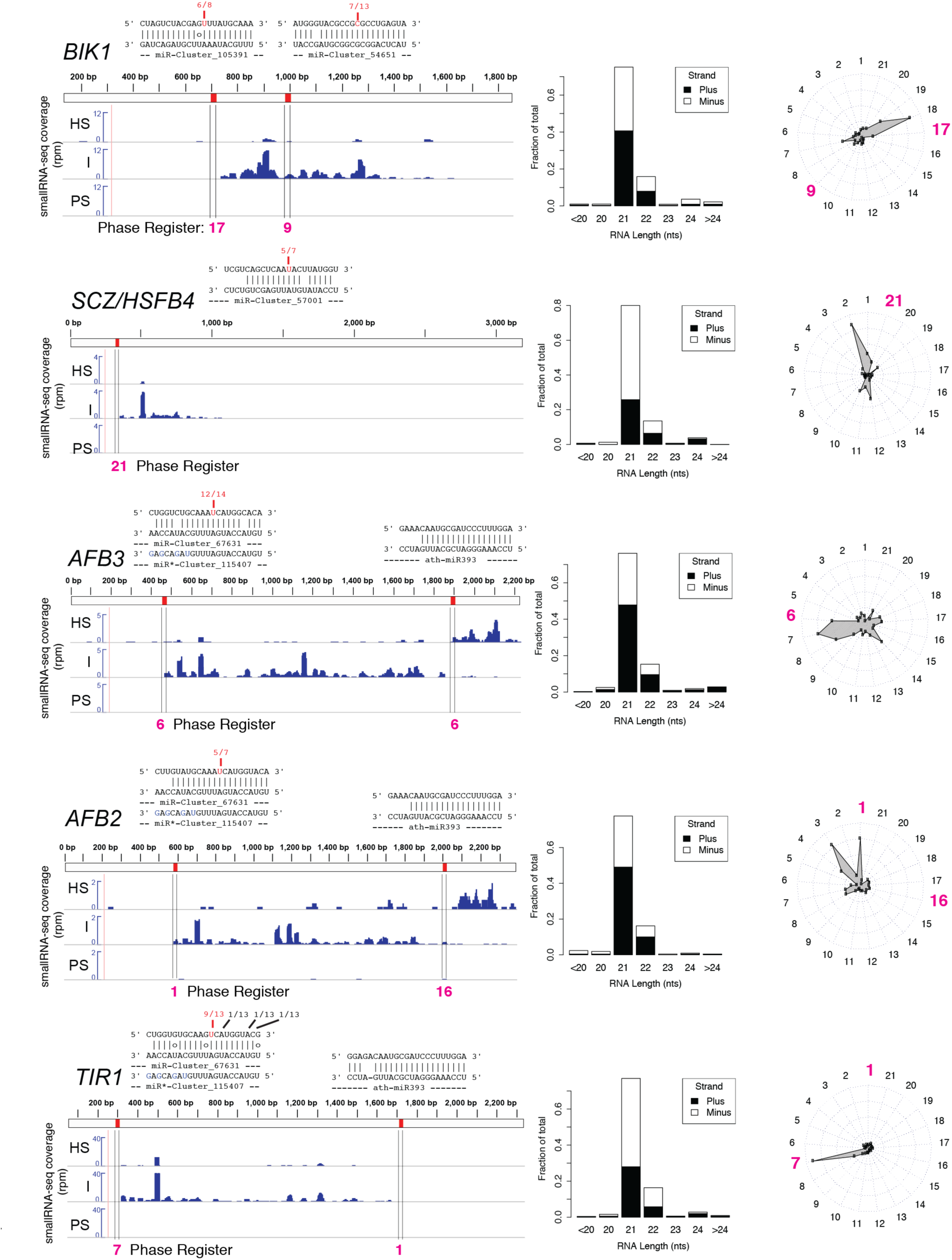
*C. campestris* miRNAs cause slicing and phased siRNA production from host mRNAs. Small RNA-seq coverage across the indicated *A. thaliana* transcripts are shown in blue for host stem (HS), interface (I), and parasite stem (PS) samples. For display, the two biological replicates of each type were merged. y-axis is in units of reads per million. Red marks and vertical lines show position of complementary sites to *C. campestris* miRNAs, with the alignments shown above. Fractions indicate numbers of 5'-RLM-RACE clones with 5'-ends at the indicated positions; the locations in red are the predicted sites for miRNA-directed slicing remnants. Bar charts show the length and polarity distribution of transcript-mapped siRNAs. Radar charts show the fractions of siRNAs in each of the 21 possible phasing registers; the registers highlighted in magenta are the ones predicted by the miRNA target sites.

**Extended Data Figure 3.**
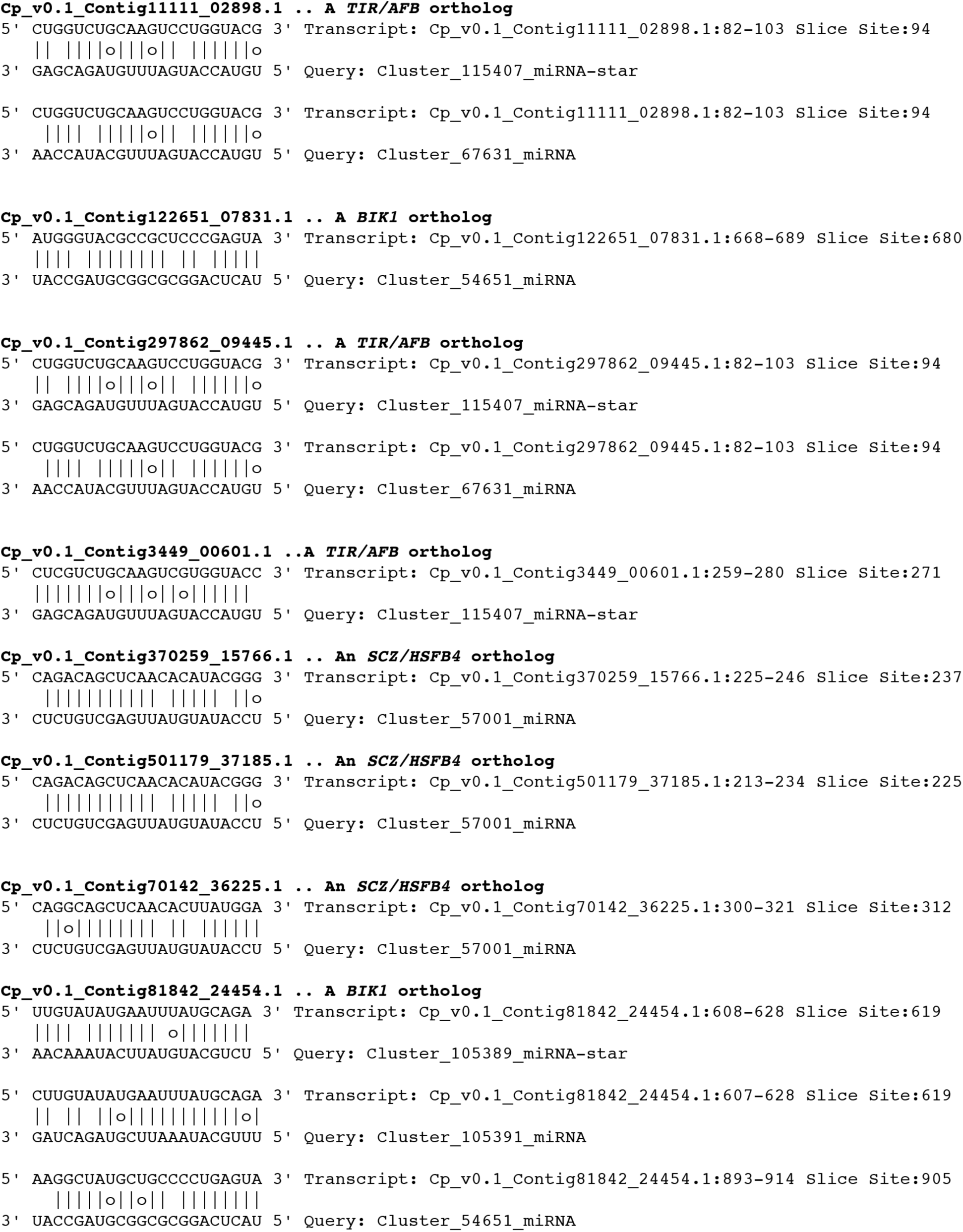
Possible miRNA target sites within endogenous *C. campestris* mRNAs. Note that none of these mRNAs had evidence of any secondary siRNA accumulation, and the complementarity of these sites was generally poor.

**Extended Data Figure 4.**
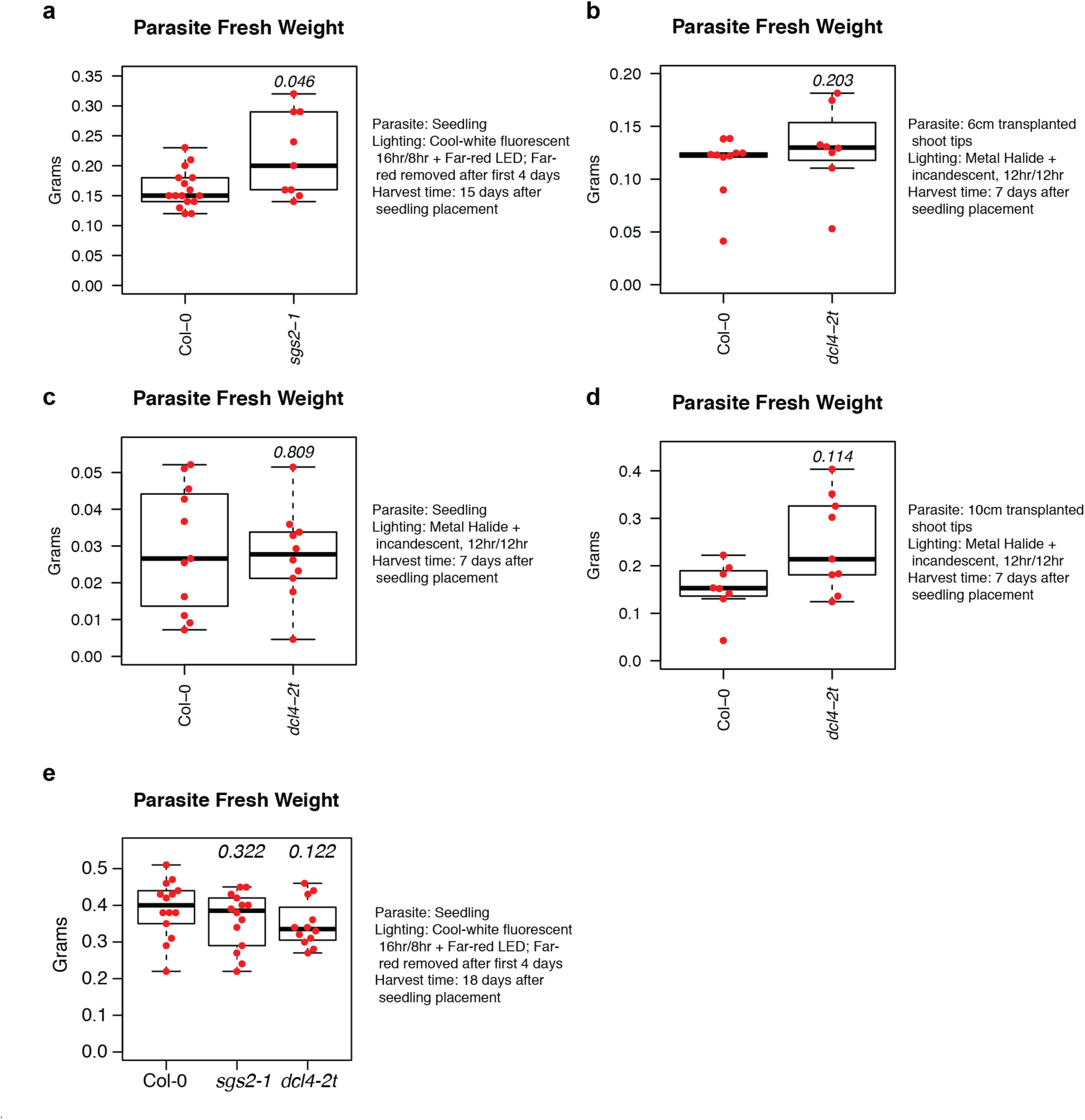
Growth of *C. campestris* on *A. thaliana sgs2-1* and *dcl4-2t* mutants with varying methodologies, as indicated. P-values (Wilcoxon rank-sum tests, unpaired, two-tailed) from comparison of mutant to wild-type (Col-0) are shown. Dots show all data points. Boxplots represent medians (horizontal lines), the central half of the data (boxes), and other data out to 1.5 times the interquartile range (whiskers). **a)** n=16 and 9 for Col-0 and *sgs2-1*, respectively. **b)** n=10 and 8 for Col-0 and *dcl4-2t*, respectively. **c)** n=11 and 10 for Col-0 and *dcl4-2t*, respectively. **d)** n=8 and 9 for Col-0 and *dcl4-2t*, respectively. **e)** n=14, 14, and 12 for Col-0, *sgs2-1*, and *dcl4-2t*, respectively.

**Extended Data Figure 5.**
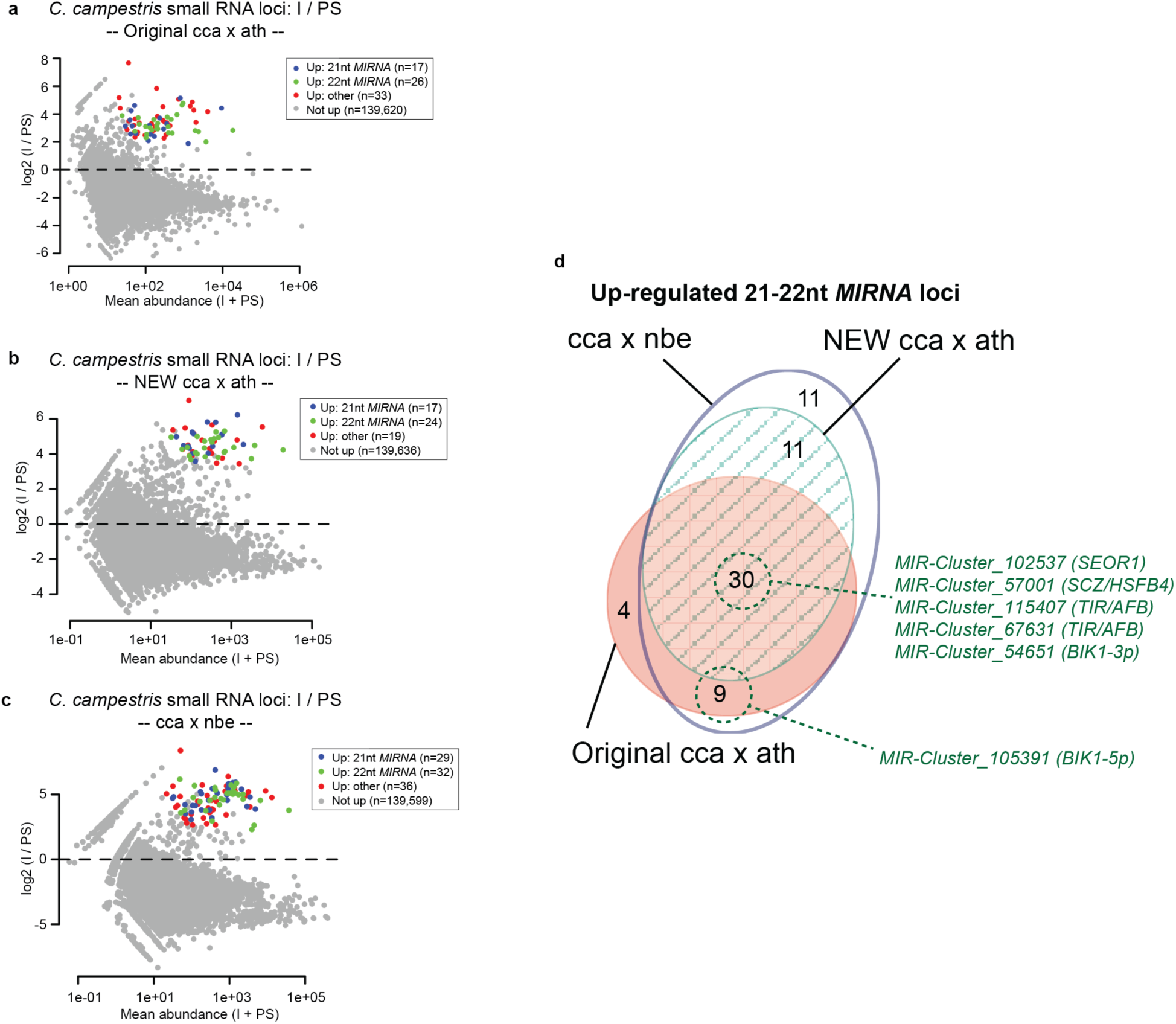
Highly reproducible induction of *C. campestris MIRNA*s on different hosts. **a)** MA plot from original experiment on *A. thaliana* hosts of *C. campestris* small RNA loci comparing interface (I) to parasite stem (PS) samples. Significantly up-regulated loci (alternative hypothesis: true difference > 2-fold, FDR <= 0.05) are highlighted. Reproduced from Figure 1A. **b-c)** As in a, except for a new set of *A. thaliana* hosts (b) or from an experiment using *Nicotiana benthamiana* as hosts (c). **d)** Area-proportional Euler diagram showing overlaps of up-regulated *C. campestris* 21-22 nt *MIRNA* loci between the three small RNA-seq experiments. The locations of the six *MIRNA* loci of special interest are highlighted in green.

**Extended Data Figure 6.**
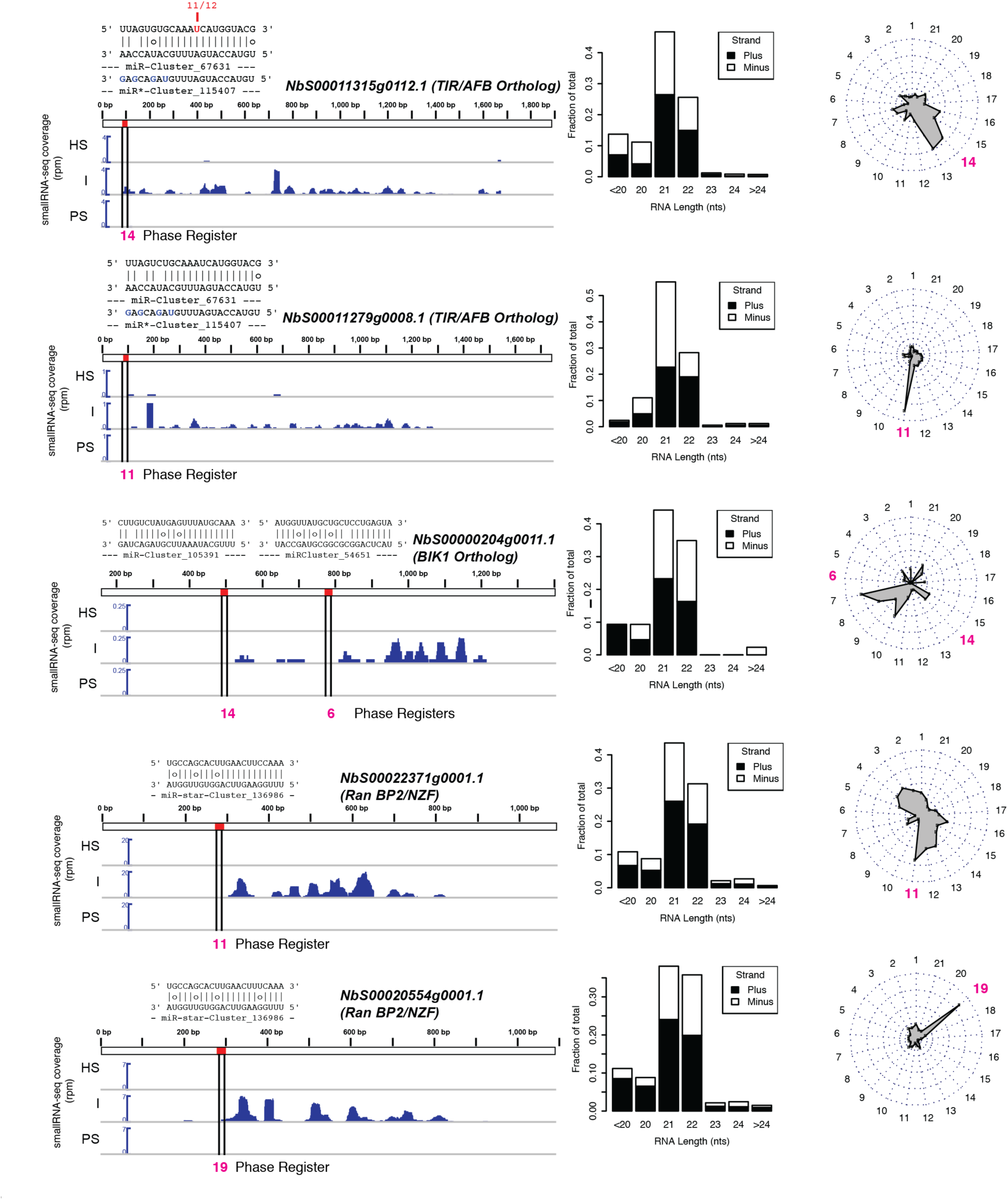
*C. campestris* miRNAs cause slicing and phased siRNA production from *Nicotiana benthamiana* mRNAs. Small RNA-seq coverage across the indicated *N. benthamiana* transcripts are shown in blue for host stem (HS), interface (I), and parasite stem (PS) samples. For display, the two biological replicates of each type were merged. y-axis is in units of reads per million. Red marks and vertical lines show position of complementary sites to *C. campestris* miRNAs, with the alignments shown above. Fraction indicates numbers of 5'-RLM-RACE clones with 5'-ends at the indicated positions; the locations in red are the predicted sites for miRNA-directed slicing remnants. Barcharts show the length and polarity distribution of transcript-mapped siRNAs. Radar charts show the fractions of siRNAs in each of the 21 possible phasing registers; the registers highlighted in magenta are the ones predicted by the miRNA target sites.

**Extended Data Figure 7.**
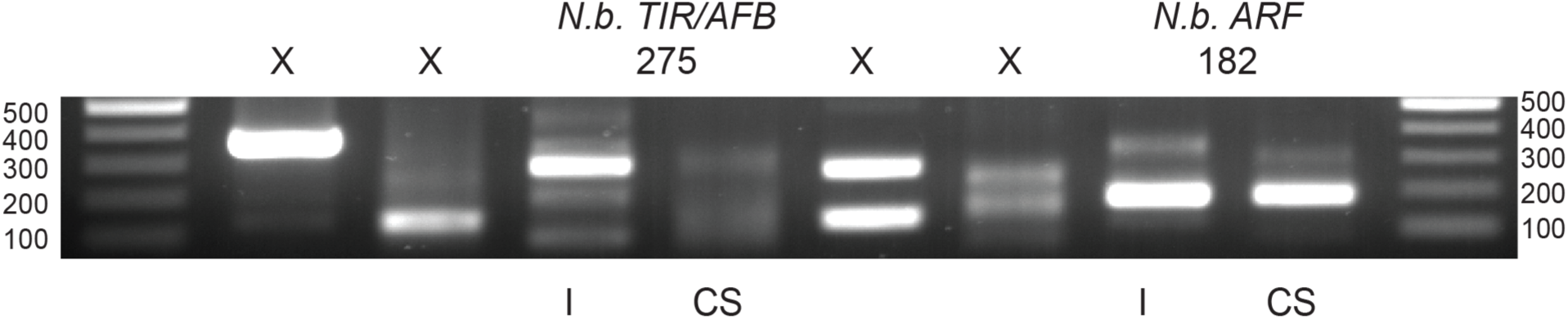
Uncropped image of *N. benthamiana* 5'-RLM-RACE products. Lanes with 'X' are irrelevant to this study. This is the uncropped version of the image in Figure 4C.

**Extended Data Figure 8.**
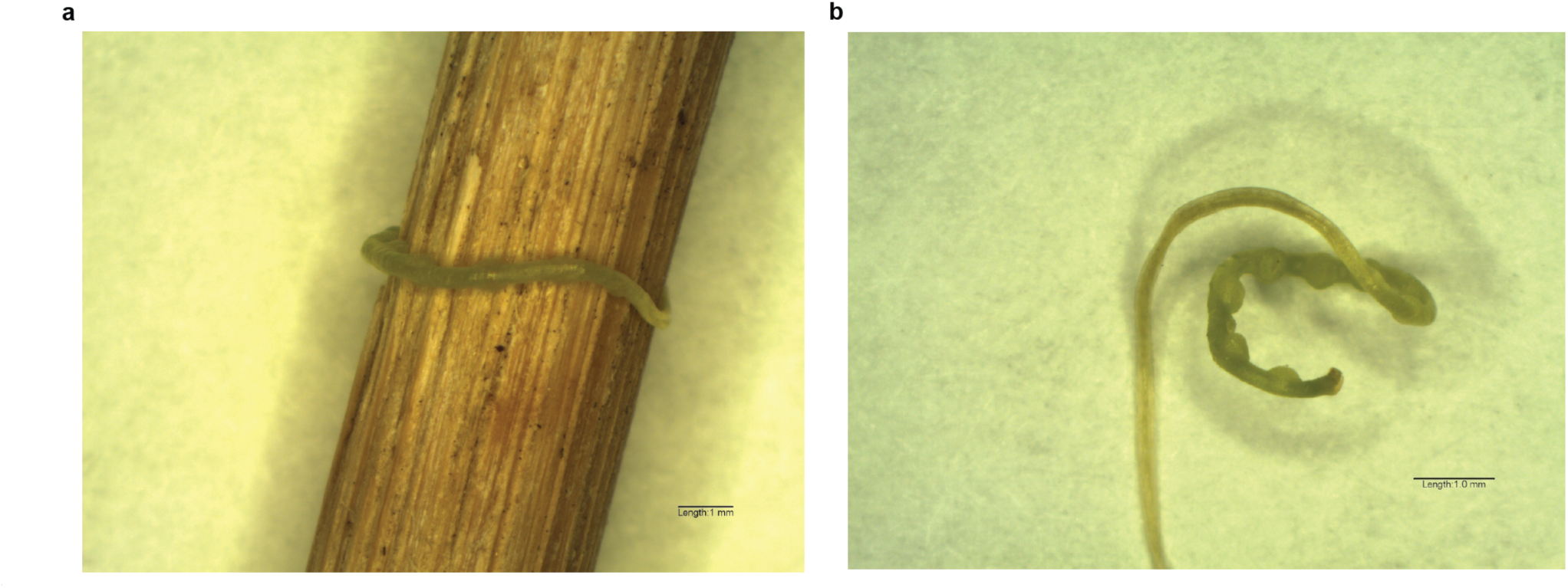
*C. campestris* pre-haustoria. **a)** *C. campestris* seedling wound around a bamboo stake. **b)** The same seedling, removed from the stake to show the prominent pre-haustorial bumps. Seedling was scarified, germinated on moist paper towels for three days at ∼28C, and then placed next to bamboo stake for four days with far-red LED lighting. Approximately 30 such seedlings were used for the 'PH' RNA in Figure 4B. Scales bars: 1mm.

**Extended Data Table 1.**
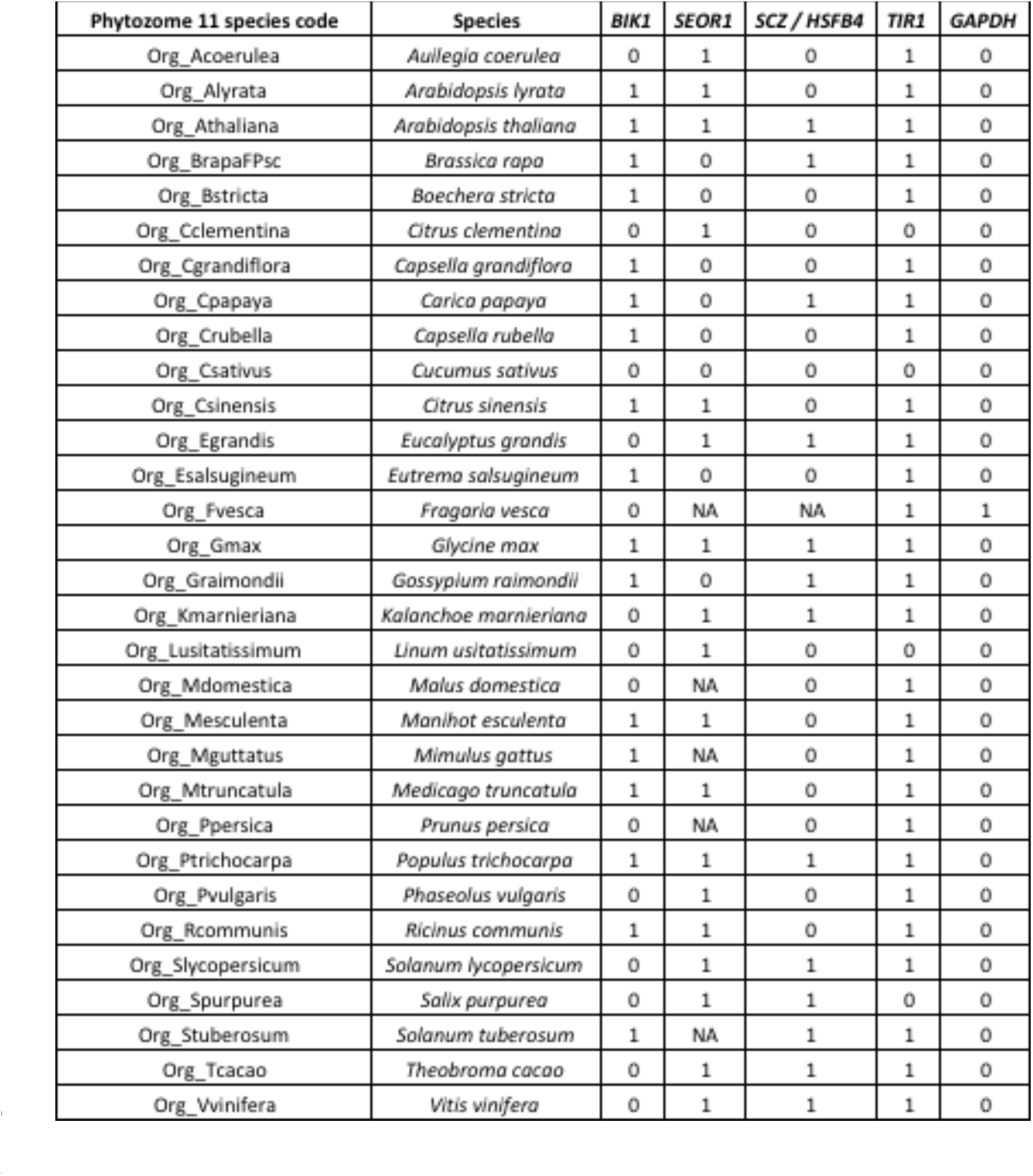
Predicted miRNA targets un multiple plant species. Targets were predicted using targetdinder.pl, keeping all hits with a score of 4.5 or less. Probable orthologs of the indicated Arabidopsis thaliana genes were found using BLASTP against the 31 eudicot species present in Phytozome 11, simply keeping up to the top 100 BLAST hits. miRNA queries were all mature miRNAs and miRNA*’s from C. campestris interface-induced MIRNA loci. An entry of 'NA' means no probable orthologs were recovered from a given species. A 1 means there was one or more predicted target in that species, a 0 means there were 0 predicted targets. GAPDH orthogroup: negative control.

| Phytozome 11 species code | Species | BIK1 | SEOR1 | SCZ / HSF4 | TIR1 | GAPDH |
| --- | --- | --- | --- | --- | --- | --- |
| Org_Acoerulea | *Auilegia coerulea* | 0 | 1 | 0 | 1 | 0 |
| Org_Alyrata | *Arabidopsis lyrata* | 1 | 1 | 0 | 1 | 0 |
| Org_Athaliana | *Arabidopsis thaliana* | 1 | 1 | 1 | 1 | 0 |
| Org_BrapaFPsc | *Brassica rapa* | 1 | 0 | 1 | 1 | 0 |
| Org_Bstricta | *Boechera stricta* | 1 | 0 | 0 | 1 | 0 |
| Org_Cclementina | *Citrus clementina* | 0 | 1 | 0 | 0 | 0 |
| Org_Cgrandiflora | *Capsella grandiflora* | 1 | 0 | 0 | 1 | 0 |
| Org_Cpapaya | *Carica papaya* | 1 | 0 | 1 | 1 | 0 |
| Org_Crubella | *Capsella rubella* | 1 | 0 | 0 | 1 | 0 |
| Org_Csativus | *Cucumis sativus* | 0 | 0 | 0 | 0 | 0 |
| Org_Csinensis | *Citrus sinensis* | 1 | 1 | 0 | 1 | 0 |
| Org_Egrandis | *Eucalyptus grandis* | 0 | 1 | 1 | 1 | 0 |
| Org_Esalsugineum | *Eutrema salsugineum* | 1 | 0 | 0 | 1 | 0 |
| Org_Fvesca | *Fragaria vesca* | 0 | NA | NA | 1 | 1 |
| Org_Gmax | *Glycine max* | 1 | 1 | 1 | 1 | 0 |
| Org_Graimondii | *Gossypium raimondii* | 1 | 0 | 1 | 1 | 0 |
| Org_Kmarnieriana | *Kalanchoe marnieriana* | 0 | 1 | 1 | 1 | 0 |
| Org_Lusitatissimum | *Linum usitatissimum* | 0 | 1 | 0 | 0 | 0 |
| Org_Mdomestica | *Malus domestica* | 0 | NA | 0 | 1 | 0 |
| Org_Mesculenta | *Manihot esculenta* | 1 | 1 | 0 | 1 | 0 |
| Org_Mguttatus | *Mimulus guttatus* | 1 | NA | 0 | 1 | 0 |
| Org_Mtruncatula | *Medicago truncatula* | 1 | 1 | 0 | 1 | 0 |
| Org_Ppersica | *Prunus persica* | 0 | NA | 0 | 1 | 0 |
| Org_Ptrichocarpa | *Populus trichocarpa* | 1 | 1 | 1 | 1 | 0 |
| Org_Pvulgaris | *Phaseolus vulgaris* | 0 | 1 | 0 | 1 | 0 |
| Org_Rcommunis | *Ricinus communis* | 1 | 1 | 0 | 1 | 0 |
| Org_Slycopersicum | *Solanum lycopersicum* | 0 | 1 | 1 | 1 | 0 |
| Org_Spurpurea | *Salix purpurea* | 0 | 1 | 1 | 0 | 0 |
| Org_Stuberosum | *Solanum tuberosum* | 1 | NA | 1 | 1 | 0 |
| Org_Tcacao | *Theobroma cacao* | 0 | 1 | 1 | 1 | 0 |
| Org_Vvinifera | *Vitis vinifera* | 0 | 1 | 1 | 1 | 0 |

## METHODS

### Germplasm

*Cuscuta* was initially obtained from a California tomato field, and seed stocks derived from self-pollination through several generations in the Westwood laboratory. The isolate was initially identified as *Cuscuta pentagona* (Engelm.) *C. pentagona* is very closely related to *C. campestris* (Yunck.), and the two are distinguished by microscopic differences in floral morphology; because of this they have often been confused^1^. We subsequently determined that our isolate is indeed *C. campestris*. *Arabidopsis thaliana sgs2-1* mutants^2^ were a gift from Hervé Vaucheret (INRA Versailles, France). *A. thaliana dcl4-2t* mutants (GABI_160G05^3^) were obtained from the Arabidopsis Biological Resource Center (The Ohio State University, USA). *A. thaliana seor* mutants (GABI-KAT 609F04^4^) were a gift from Michael Knoblauch (Washington State University, USA). The *A. thaliana tir1-1/afb2-* and *afb3-*4 mutants^5^ were a gift from Gabriele Monshausen (The Pennsylvania State University, USA). The *bik1* mutant^6^ was a gift from Tesfaye Mengiste (Purdue University, USA). The *scz2* mutant^7^ was a gift from Renze Heidstra (Wageningen University, The Netherlands). All *A. thaliana* mutants were in the Col-0 background.

### Growth conditions and RNA extractions

For initial experiments (small RNA-seq and RNA blots in Figure 1) *A. thaliana* (Col-0) plants were grown in a growth room at 18-20°C with 12-h light per day, illuminated (200 μmol m^-2^s^-1^) with metal halide (400W, GE multi-vapor lamp) and spot-gro (65W, Sylvania) lamps. *C. campestris* seeds were scarified in concentrated sulfuric acid for 45 min, followed by 5-6 rinses with distilled water and dried. *C. campestris* seeds were placed in potting medium at the base of four-week-old *A. thaliana* seedlings and allowed to germinate and attach to hosts. The *C. campestris* plants were allowed to grow and spread on host plants for an additional three weeks to generate a supply of uniform shoots for use in the experiment. Sections of *C. campestris* shoot tip (∼10 cm long) were placed on the floral stem of a fresh set of *A. thaliana* plants. Parasite shoots coiled around the host stems and formed haustorial connections. Tissues from plants that had established *C. campestris* with at least two coils around healthy host stems and clear parasite growth were used in these studies. Control plants were grown under the same conditions as parasitized plants, but were not exposed to *C. campestris*.

For the preparation of tissue-specific small RNA libraries, tissues were harvested after *C. campestris* cuttings had formed active haustorial connections to the host. This was evidenced by growth of the *C. campestris* shoot to a length of at least 10 cm beyond the region of host attachment (7-10 d after infection). Three tissues were harvested from the *A. thaliana*-*C. campestris* associations: 1) 2.5 cm of *A. thaliana* stem above the region of attachment, 2) *A. thaliana* and *C. campestris* stems in the region of attachment (referred to as the interface), 3) 2.5 cm of the parasite stem near the point of attachment. To remove any possible cross-contamination between *A. thaliana* and *C. campestris*, harvested regions of the parasite and host stem were taken 1 cm away from the interface region and each harvested tissue was surface cleaned by immersion for 5 min in 70% ethanol, the ethanol was decanted and replaced, the process was repeated three times and the stems were blotted dry with a Kimwipe after the final rinse. All three sections of tissue were harvested at the same time and material from 20 attachments were pooled for small RNA extraction. Small RNA was extracted from ∼ 100 mg of each tissue using the mirPremier microRNA Isolation Kit (Sigma-Aldrich, St. Louis, MO, USA) according to the manufacturer’s protocol. Small RNA was analyzed using an Agilent small RNA Kit on a 2100 Bioanalyzer platform.

Samples used for RNA ligase-mediated 5' rapid amplification of cDNA ends (5'-RLM-RACE; Figure 2D), quantitative reverse-transcriptase polymerase chain reaction (qRT-PCR; Figure 3A) analyses of *A. thaliana* targets were prepared as described above with the following modifications: Col-0 *A. thaliana* hosts were cultivated in a growth room with 16 hr. days, 8 hr. nights, at ∼23C, under cool-white fluorescent lamps, attachment of *C. campestris* cuttings was promoted by illumination with far-red LED lighting for 3-5 days, and total RNA was extracted using Tri-reagent (Sigma) per the manufacturer’s suggestions, followed by a second sodium-acetate / ethanol precipitation and wash step. Samples used for RNA blots of secondary siRNA accumulation from *A. thaliana* mutants and replicate small RNA-seq libraries were obtained similarly, except that the samples derived from the primary attachments of *C. campestris* seedlings on the hosts instead of from cuttings. In these experiments, scarified *C. campestris* seedlings were first germinated on moistened paper towels for three days at ∼28C, then placed adjacent to the host plants with their radicles submerged in a water-filled 0.125ml tube.

*C. campestris* pre-haustoria (Extended Data Figure 8) were obtained by scarifying, germinating and placing seedlings as described above, next to bamboo stakes in soil, under illumination from cool-white fluorescent lights and far-red emitting LEDs. Seedlings coiled and produced pre-haustoria four days after being placed, and were harvested and used for total RNA extraction (used for RNA blot in Figure 4B) using Tri-reagent as described above. *Nicotiana benthamiana* was grown in a growth room with 16 hr. days, 8 hr. nights, at ∼23C, under cool-white fluorescent lamps. Three-to four-week old plants served as hosts for scarified and germinated *C. campestris* seedlings. Attachments were promoted by three-six days with supplementation by far-red emitting LEDs. Under these conditions, *C. campestris* attached to the petioles of the *N. benthamiana* hosts, not the stems. Interfaces and control petioles from un-parasitized hosts were collected 7-8 days after successful attachments, and total RNA (used for RNA blots in Figure 4B and small RNA-seq libraries) recovered using Tri-reagent as described above.

### small RNA-seq

The initial small RNA-seq libraries were constructed using the Illumina Tru-Seq small RNA kit per the manufacturer’s protocol and sequenced on an Illumina HiSeq2500 instrument. Subsequent small RNA-seq libraries (replicate two using *A. thaliana* hosts, and the *N. benthamiana* experiments) instead used New England Biolabs NEBnext small RNA library kit, following the manufacturer’s instructions. Raw sRNA-seq reads were trimmed to remove 3'-adapters, and filtered for quality and trimmed length >= 16 nts using cutadapt^8^ version 1.9.1 with settings "-a TGGAATTCTCGGGTGCCAAGG –discard-untrimmed -m 16 –max-n=0". For experiments where *A. thaliana* was the host, trimmed reads that aligned with zero or one mismatch (using bowtie^9^ version 1.1.2, settings "-v 1") to the *A. thaliana* plastid genome, the *C. gronovii* plastid genome (*C. gronovii* was the closest relative to *C. campestris* that had a publically available completed plastid genome assembly available), *A. thaliana* rRNAs, tRNAs, snRNAs, or snoRNAs were removed. Similarly, for experiments where *N. benthamiana* was the host, the reads were cleaned against the *C. gronovii* plastid genome, the *N. tabacum* plastid genome and rRNAs, and a set of tRNAs predicted from the *N. benthamiana* genome using tRNAscanSE.

For the original *A. thaliana* host data, the 'clean' reads were aligned and analyzed with reference to the combined TAIR10 *A. thaliana* reference genome and a preliminary version 0.1 draft genome assembly of *C. campestris* using ShortStack^10^ (version 3.8.3) using default settings. The resulting annotated small RNA loci (Supplementary Data 1) were analyzed for differential expression (I vs. PS) using DESeq2^11^, with a log_2_ fold threshold of 1, alternative hypothesis of "greaterAbs", and alpha of 0.05. p-values were adjusted for multiple testing using the Benjamini-Hochberg procedure, and loci with an adjusted p-value of <= 0.05 (equivalent to an FDR of <= 0.05) were called up-regulated in I relative to PS. Among the up-regulated loci, those annotated by ShortStack as *MIRNA*s deriving from the *C. campestris* genome which produced either a 21nt or 22nt mature miRNA (Supplementary Data 2) were retained and further analyzed. The predicted secondary structures and observed small RNA-seq read coverage was visualized (Supplementary Data 3-4) using strucVis (version 0.3; <u>https://github.com/MikeAxtell/strucVis</u>).

For analysis of mRNA-derived secondary siRNAs, the 'clean' small RNA-seq reads from the original *A. thaliana* experiment were aligned to the combined TAIR10 representative cDNAs from *A. thaliana* and our preliminary version 0.1 transcriptome assembly for *C. campestris*, using ShortStack^10^ version 3.8.3, with settings –mismatches 0, –nohp, and defining the full length of each mRNA as a 'locus' using option –locifile. The resulting counts of small RNA alignments for each mRNA were used for differential expression analysis, comparing I vs. HS, using DESeq2 ^11^ as described above. *A. thaliana* mRNAs with significantly up-regulated (FDR <= 0.05) small RNAs comparing I vs. HS were retained for further analysis. The cDNA sequences of these loci were retrieved, and used for miRNA target predictions using GSTAr (version 1.0; <u>https://github.com/MikeAxtell/GSTAr</u>); the full set of mature miRNAs and miRNA*’s (Supplementary Data 2) from the I-induced *C. campestris MIRNA* loci were used as queries.

Analysis of the second set of *A. thaliana* - *C. campestris* small RNA-seq data aligned the cleaned reads to the combined *A. thaliana* and *C. campestris* reference genomes as described above, except that the list of loci derived in the analysis of the original data (Supplementary Data 1) was used as a "-locifile" in the ShortStack analysis. Differential expression analysis was then performed using DESeq2 as described above. Analysis of the *N. benthamiana - C. campestris* small RNA-seq data began with a ShortStack analysis of the cleaned reads against the combined *N. benthamiana* (version 0.4.4) genome and the preliminary assembly of the *C. campestris* genome, using default settings. The *de novo N. benthamiana* loci obtained from this run were retained. The resulting alignments were used to quantify small RNA abundance from the *C. campestris* small RNA loci defined with the original data. The resulting read counts were then used for differential expression analysis with DESeq2 as described above. Analysis of secondary siRNAs derived from *N. benthamiana* mRNAs was performed similarly as the *A. thaliana* mRNA analysis described above, except that the combined transcriptomes were from *C. campestris* and *N. benthamiana* (version 0.4.4 annotations).

### RNA blots

Small RNA gel blots were performed as previously described^12^ with modifications. For the blots in Figure 1B, small RNAs (1.8 micrograms) from each sample were separated on 15% TBE-Urea Precast gels (Bio-Rad), transblotted onto the Hybond NX membrane and cross-linked using 1-ethyl-3-(3-dimethylamonipropyl) carbodiimide^13^. Hybridization was carried out in 5×SSC, 2×Denhardt’s Solution, 20 mM sodium phosphate (pH 7.2), 7% SDS with 100 μg/ml salmon testes DNA (Sigma-Aldrich). Probe labeling, hybridization and washing were performed as described^12^. Radioactive signals were detected using Typhoon FLA 7000 (GE Healthcare). Membranes were stripped in between hybridizations by washing with 1% SDS for 15 min at 80°C and exposed for at least 24 h to verify complete removal of probe before re-hybridization. Sequences of probes are listed below. Blots in Figures 3B and 4B were performed similarly, except that 12 micrograms of total RNA were used instead. Probe sequences are listed in Supplementary Data 6.

### 5' RNA ligase-mediated rapid amplification of cDNA ends (5'-RLM-RACE)

Five micrograms of total RNA were ligated to one microgram of a 44 nucleotide RNA adapter (Supplementary Data 6) using a 20ul T4 RNA ligase 1 reaction (NEB) per the manufacturer’s instructions for a one-hour incubation at 37C. The reaction was then diluted with 68ul of water and 2ul of 0.5M EDTA pH 8.0, and incubated at 65C for 15 minutes to inactivate the ligase. Sodium acetate pH5.2 was added to a final concentration of 0.3M, and the RNA precipitated with ethanol. The precipitated and washed RNA was resuspended in 10ul of water. 3.33ul of this sample was used as the template in a reverse transcription reaction using random primers and the Protoscript II reverse transcriptase (NEB) per the manufacturer’s instructions. The resulting cDNA was used as template in first round PCR using a 5' primer matching the RNA adapter and a 3' gene-specific primer (Supplementary Data 6). 1ul of the first round PCR product was used as the template for nested PCR with nested primers (Supplementary Data 6). Gene-specific primers for *A. thaliana* cDNAs were based on the representative TAIR10 transcript models, while those for *N. benthamiana* cDNAs were based on the version 0.4.4 transcripts (Sol Genomics Network^14^). In Figure 4c, *N. benthamiana TIR/AFB* is transcript ID *NbS00011315g0112.1*; *N. benthamiana ARF* is transcript ID *NbS00059497g0003.1*. Bands were gel-purified from agarose gels and cloned into pCR4-TOPO (Life Tech). Inserts from individual clones were recovered by colony PCR and subject to Sanger sequencing.

### Quantitative reverse-transcriptase PCR (qRT-PCR)

Total RNA used for qRT-PCR was first treated with DNaseI (RNase-free; NEB) per the manufacturer’s instructions, ethanol precipitated, and resuspended. 2 micrograms of treated total RNA was used for cDNA synthesis using the High Capacity cDNA Synthesis Kit (Applied Biosystems) per the manufacturer’s instructions. PCR reactions used PerfeCTa SYBR Green FastMix (Quanta bio) on an Applied Biosystems StepONE-Plus quantitative PCR system per the manufacturer’s instructions. Primers (Supplementary Data 6) were designed to span the miRNA target sites, to ensure that only uncleaved mRNAs were measured. Three reference mRNAs were used: *ACTIN*, *PP2A (PP2A sub-unit PDF2*; *At1g13320*), and *TIP41-l* (*TIP41-like; At4g34270*) ^15^. Raw Ct values were used to calculate relative normalized expression values to each reference mRNA separately, and the final analysis took the median relative expression values between the *ACTIN-* and *TIP41-l* normalized data.

### *C. campestris* growth assays

*C. campestris* seedlings were scarified, pre-germinated, and placed next to hosts in 0.125ml water-filled tubes under cool-white fluorescent lighting supplemented with far-red emitting LEDs (16hr day, 8hr night) at ∼ 23C as described above. After a single attachment formed (4 days), far-red light supplementation was removed to prevent secondary attachments. After 18 more days of growth, entire *C. campestris* vines were removed and weighed (Figure 3C). Multiple additional growth trials were performed specifically on the *dcl4-2t* and *sgs2-1* mutant hosts under varying conditions (Extended Data Figure 4).

### miRNA target predictions

To find probable orthologs for *Arabidopsis thaliana* genes of interest, the *A. thaliana* protein sequences were used as queries for a BLASTP analysis of the 31 eudicot proteomes available on Phytozome 11 (<u>https://phytozome.jgi.doe.gov/pz/portal.html#</u>). Transcript sequences for the top 100 hits were retrieved. In some cases no hits from a particular species were found; these are 'N/A' on Figure 4a. The miRNA query set was all mature miRNAs and miRNA*s from the I-induced, *C. campestris*-derived 21nt or 22nt *MIRNA*s (Supplementary Data 2). Targets were predicted from the probable 31-species with a maximum score of 4.5 using targetfinder.pl (<u>https://github.com/MikeAxtell/TargetFinder/</u>) version 0.1.

*N. benthamiana* orthologs of *A. thaliana* proteins were found based on BLAST-P searches against the version 0.4.4 *N. benthamiana* protein models at Sol Genomics Network^14^, and miRNA target sites predicted using targetfinder.pl as above.

### Code availability

ShortStack^10^ (small RNA-seq analysis), strucVis (visualization of predicted RNA secondary structures with overlaid small RNA-seq depths), and Shuffler.pl/targetfinder.pl (prediction of miRNA targets controlling for false discovery rate) are all freely available at <u>https://github.com/MikeAxtell</u>. Cutadapt version 1.9.1^8^ is freely available at http://cutadapt.readthedocs.io/en/stable/index.html. The R package DESeq2^11^ is freely available at http://www.bioconductor.org/packages/release/bioc/html/DESeq2.html.

### Data Availability

Small RNA-seq data from this work are available at NCBI GEO under accession GSE84955 and NCBI SRA under project PRJNA408115. The draft, preliminary *C. campestris* genome and transcriptome assemblies used in this study are available at the Parasitic Plant Genome Project website at http://ppgp.huck.psu.edu.

## SI Guide

**Supplementary Data 1:** *A. thaliana* and *C. campestris* small RNA loci. Output from ShortStack 3.8.3 (<u>https://github.com/MikeAxtell/ShortStack</u>) showing all small RNA loci identified in this study. (.xlsx format)

**Supplementary Data 2**: Mature miRNAs and miRNA*s from *C. campestris* Interface-induced *MIRNA* loci. (.xlsx format)

**Supplementary Data 3**: Details of *C. campestris MIRNA* loci: Text-based sequences, predicted secondary structures, and aligned small RNA reads (all six libraries). Lower-case letters indicate small RNA bases that are mismatched to the genomic sequence. Plain-text (ASCII) format.

**Supplementary Data 4**: Images showing *C. campestris* hairpins overlaid with color-codes representing total read-depth (all six 'original' *C. campestris* x *A. thaliana* small RNA libraries). (pdf, 43 pages).

**Supplementary Data 5**: Alignments of mature miRNAs and miRNA*s (from Supplementary Data 2 – the induced *C. campestris* miRNAs) against the *A. thaliana* genome. Note that most have three or more mismatches. (SAM format).

**Supplementary Data 6**: Oligonucleotide sequences. Excel (.xlsx) format.

